# Morphodynamical cell state description via live-cell imaging trajectory embedding

**DOI:** 10.1101/2021.10.07.463498

**Authors:** Jeremy Copperman, Sean M. Gross, Young Hwan Chang, Laura M. Heiser, Daniel M. Zuckerman

## Abstract

Time-lapse imaging is a powerful approach to gain insight into the dynamic responses of cells, but the quantitative analysis of morphological changes over time remains challenging. Here, we exploit the concept of “trajectory embedding” to analyze cellular behavior using morphological feature trajectory histories—that is, multiple time points simultaneously, rather than the more common practice of examining morphological feature time courses in single timepoint (snapshot) morphological features. We apply this approach to analyze live-cell images of MCF10A mammary epithelial cells after treatment with a panel of microenvironmental perturbagens that strongly modulate cell motility, morphology, and cell cycle behavior. Our morphodynamical trajectory embedding analysis constructs a shared cell state landscape revealing ligand-specific regulation of cell state transitions and enables quantitative and descriptive models of single-cell trajectories. Additionally, we show that incorporation of trajectories into single-cell morphological analysis enables (i) systematic characterization of cell state trajectories, (ii) better separation of phenotypes, and (iii) more descriptive models of ligand-induced differences as compared to snapshot-based analysis. This morphodynamical trajectory embedding is broadly applicable to the quantitative analysis of cell responses via live-cell imaging across many biological and biomedical applications.

## Introduction

In normal and diseased tissues, cells are continually exposed to a wide variety of extracellular stimuli, including growth factors and cytokines, that modulate morphological and phenotypic states However, quantitative assessment of complex morphological states remains a challenging problem. Single timepoint measurements provide some information about cell state but do not capture how responses evolve over time. Live-cell imaging has been deployed to characterize dynamical changes in cell morphology, or cellular morphodynamics^1, 2^. Recent advances in live-cell imaging technologies have enabled unprecedented assessment of the behavior and interactions of cellular populations^3–6^. To date, however, most analyses of live-cell image data have been primarily based upon classification of cell morphology observed in individual time points and do not directly examine the rich dynamic landscape of cell morphology trajectories^7–9^.

Here we describe a generalizable morphodynamical trajectory embedding method to analyze live-cell imaging datasets composed of unlabeled phase-contrast microscopy images. Analysis methods directly based upon trajectory features that are aggregated over multiple time points have been used to classify mitosis and apoptosis^10^, and also to monitor signaling responses via reporter molecules^11^. We show here that by mapping the multiple-time-point *trajectory space* of cells, rather than examining single-cell time courses built from snapshots, we can increase the information extracted from live-cell imaging experiments and improve the quantitative description of cellular responses.

Live-cell imaging provides temporal information not available from other single-cell and omic measures. Single-cell RNA sequencing can assay thousands of molecular read-outs across thousands or hundreds of thousands of cells. A common data analysis procedure is the extraction of cell states, including continuous low-dimensional cell state spaces, from high-dimensional molecular data^12^. Because sequencing is a destructive readout, single-cell trajectories in this space can only be inferred indirectly through population time-series modeling^13–16^ or pseudo-time approaches^17, 18^; however, approaches such as “RNA velocity”^19, 20^ have been used to infer cell state dynamics. In contrast, live-cell imaging enables cellular and phenotypic responses to be assayed over multiple time points.

In a live-cell imaging experiment, cell responses to a perturbation, such as the addition of a signaling ligand to the cell culture medium, can be examined. Cells with shared response patterns can be assigned to distinct cell states that can then be tracked over time to quantify response dynamics. Cell states can be discrete, as for cell-cycle states G0-G1-G2-M, or continuous as for epithelial-to-mesenchymal transition.. Live-cell imaging has been used to develop gene-level functional annotation, including in RNAi gene knockout screens and drug screening^7^, and also to study how single-cell trajectories evolve in a cell state space that represents distinct cell-cycle phases^8^. Gordonov et al. developed an unsupervised approach to characterize live-cell state, analyze cell shape space, and obtain models of cell responses that included three distinct cell states^21^. This workflow of live-cell imaging, segmentation, featurization, and tracking has been used to describe cell state as a continuum^22^ and to develop a cell trajectory-based description of an epithelial-mesenchymal transition (EMT)^23^ in the space of single-timepoint snapshot features. Heryanto et al. utilized 3D shape descriptors to explore the relationship between 3D shape and cell motility^24^. Notably, trajectory information, including combined motility and morphological features computed as averages over single-cell trajectories, have been used to define and identify cell state space in microglia and neural progenitor cells.^25^

Trajectories are the natural space from which to classify a system out of equilibrium, such as a living cell^26, 27^. Leveraging such a framework, however, requires that the state space of the system can be measured. Floris Takens’ seminal trajectory embedding theorem^28^ states that in a deterministic dynamical system, there is a 1:1 correspondence between the space of the full dynamical system and that formed by concatenating incomplete observations of the system across time—the “trajectory embedding” space, also referred to as delay-embedding. For cell morphodynamics, the incomplete observations are single-cell morphological features (snapshots) and concatenating features across time forms morphodynamical feature trajectories, which we refer to as “trajectories.” For *N_f_* features and *n_τ_* trajectory timepoints, the trajectory of a cell can be considered a vector of dimensionality *N_f_* x *n_τ_*. In stochastic systems or systems of sufficient complexity, such as a cell, a comprehensive map enabling perfect prediction of the dynamical system is not necessarily achievable, but trajectory embedding can still lead to an improved characterization of the dynamical behavior^29–31^. Trajectory embedding methodology has been applied in fields as diverse as weather prediction^32^, economics^33^, and molecular dynamics^34–36^, but to our knowledge has not yet been applied to examine cell states observable in live-cell imaging assays.

Here, we develop and apply the morphodynamical trajectory embedding method in a dataset of MCF10A mammary epithelial cells perturbed with a set of six ligands that span major extracellular signaling pathways and induce distinct cellular responses, including changes to cell proliferation, differentiation state, and motility and which are known to be important in mammary tissues^37^. The live-cell imaging data are part of a broader data collection effort through the Library of Integrated Network-Based Cellular Signatures (LINCS) consortium^38, 39^ MCF10A project^37^ where the molecular and phenotypic responses to these ligand perturbations were assessed. Molecular and cellular responses indicate changes in multiple pathways and initiation of unique cellular responses in each ligand condition^37^. Our live-cell imaging cell-trajectory-based analysis was developed to characterize the morphodynamical changes associated with molecular responses. By directly analyzing morphological feature trajectories, rather than single-timepoint features or averaged features over time, we leverage the additional information contained in the time-ordered single-cell trajectory information.

## Results

We applied our trajectory embedding analysis to systematically characterize cell state from live-cell imaging of MCF10A mammary epithelial cells treated with a panel of ligands that induce distinct phenotypic responses: PBS (no ligand control), EGF, HGF, OSM, BMP2 + EGF, IFNG + EGF, and TGFB + EGF. Cells were assessed via phase-contrast microscopy over 48 hours, with images collected every 30 min, as part of the LINCS MCF10A project^37^. Single-cells were segmented, featurized, and tracked through time as described in Methods. In our “trajectory embedding” approach, time-sequences of features for the trajectory length under consideration were concatenated and used for UMAP^40^ dimensionality reduction as the basis for further analysis. The single-cell trajectories are the set of extracted single-cells and their tracks, or linkages between frames. An overview of the live-cell imaging trajectory embedding workflow is shown in Figure 1. The quantification of morphodynamical cell states demonstrates the biological information intrinsic to cellular trajectories in a broadly applicable and relatively simple imaging assay.

**Figure 1:**
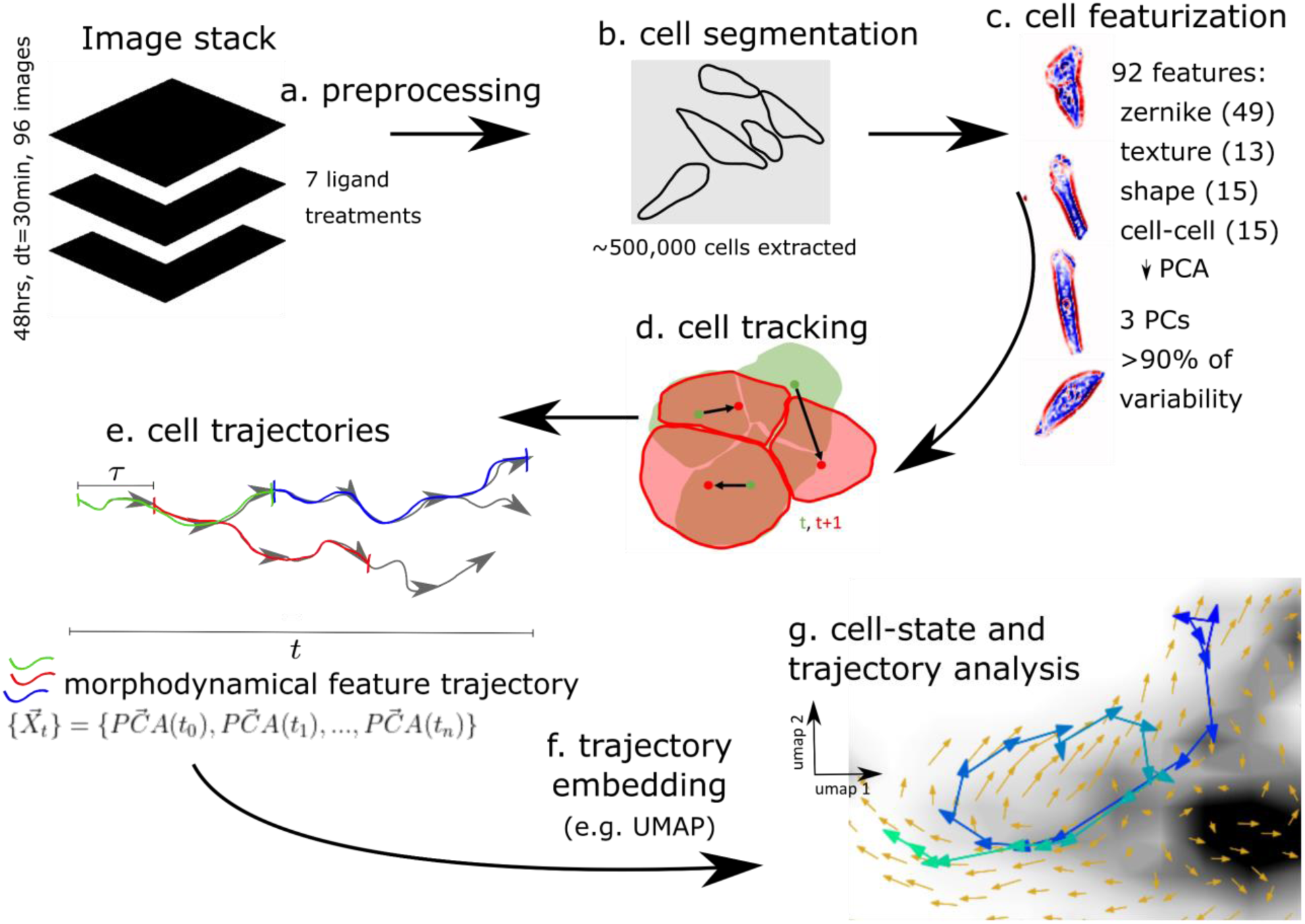
Live-cell imaging analysis and trajectory embedding pipeline. The data analysis pipeline starts from 48-hour image stacks and proceeds to the morphodynamical trajectory analysis. Image processing steps include a. preprocessing, b. cell segmentation, c. featurization (z-normalized phase-contrast pixel values colored red positive to blue negative), d. tracking (cell boundaries at t, t+30 min with cell centers connected by black arrows), e. extracting morphodynamical trajectories as sliding window cell feature trajectory snippets 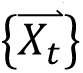 from cell linkages (3 possible trajectory snippets of length 3 shown in green, red, and blue), f. trajectory embedding (UMAP), and g. cell state and trajectory analysis in the trajectory embedding (UMAP) space using trajectories longer than the trajectory embedding length.

Comparing morphodynamical trajectories between the different ligand treatments requires the construction of a shared cell state space, which we created by analyzing all of the trajectories from the full set of treatments through the dimensionality reduction pipeline together. The single-cell trajectories we use for a trajectory embedding of length *n_τ_* consist of all possible trajectory snippets of length *n_τ_* in the full trajectory set; for example, a single cell that is tracked over 12 frames will have 5 possible trajectory snippets of length 8 in a sliding window manner (frames 1-8, 2-9, 3-10, 4-11, 5-12). Unless otherwise labeled, we used all available trajectory snippets over the 48-hour experiment. Snippets mapped to the same location in the reduced-dimensionality cell state space share qualitatively similar morphologies across trajectory timepoints and across all treatments (supplementary figure 2). In this shared morphodynamical trajectory space, ligand treatments alter the distribution of morphodynamical cell states (Figure 2).

**Figure 2:**
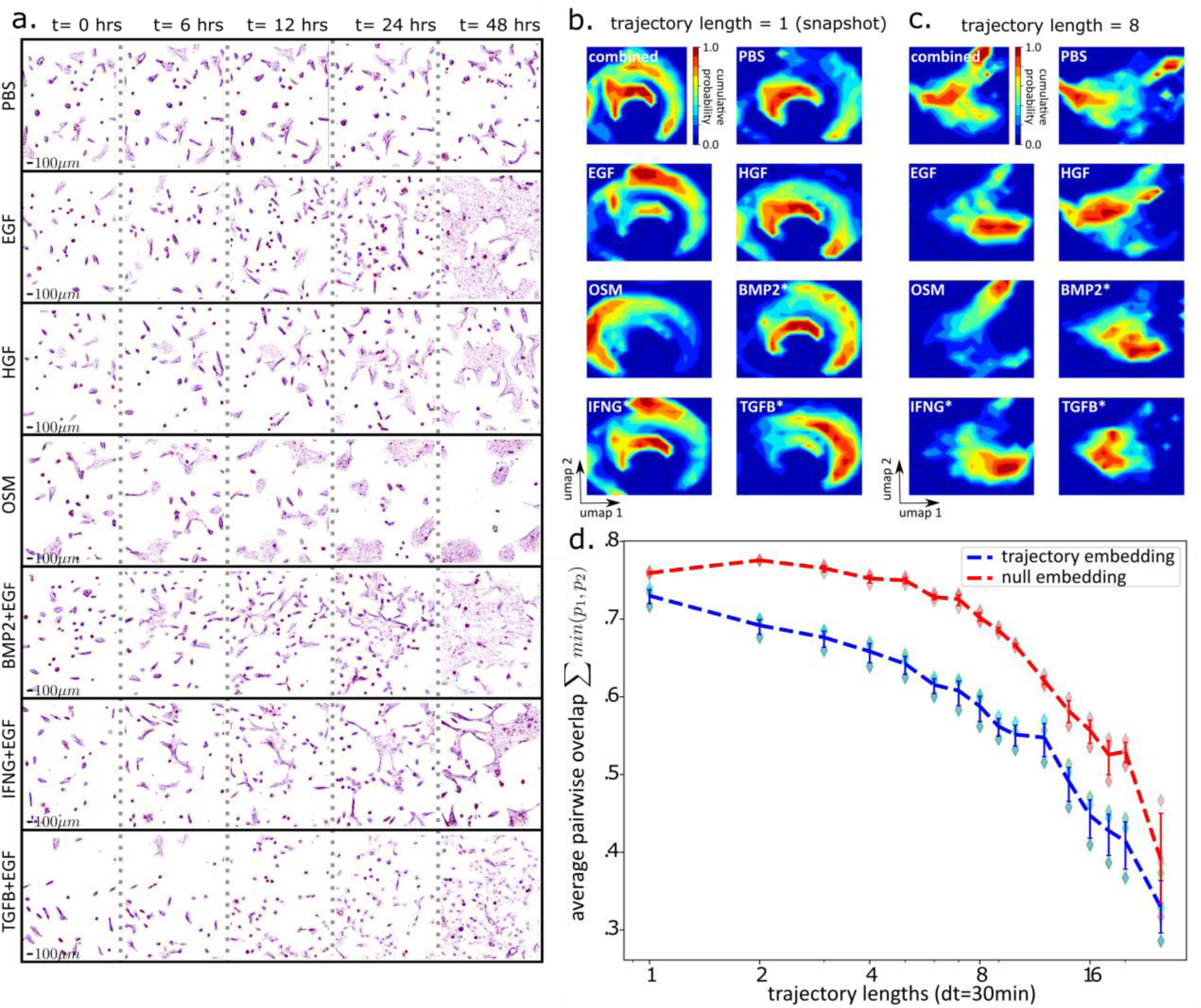
Trajectory embedding increases the distinguishability of cell states induced by ligand perturbation. a) Representative background subtracted phase-contrast images (size 1.6mm x 1.9mm with z-normalized phase-contrast pixel values colored red (positive phase contrast) to blue (negative phase contrast) in the set of ligand conditions (top to bottom) at 0, 6, 12, 24, and 48 hrs (left to right). b,c) Distributions of cells in trajectory embedding space over the 48 hours of imaging. b) snapshot space (trajectory embedding length = 1), and c) trajectory embedding length = 8. d) Average pairwise overlap over all treatment pairs (shared area under probability distributions) as a function of the morphodynamical feature trajectory length used in the embedding (log_2_ x-axis scale), comparing trajectory embedding (blue dashed lines) and null model with randomly scrambled time labels within treatments (red dashed lines), averaging over results obtained by dividing data into three sets by field of view (diamonds), with error bars from a bootstrapped 95% confidence interval over the three data splits. A trajectory length of 1 corresponds to a snapshot description.

### Separation of unique and shared cell state under ligand perturbation

Ligand perturbation can induce time-dependent morphologic and phenotypic changes, including cycling rate, motility, and cytoskeletal features. Cells in the control condition (PBS, no ligand) did not proliferate, while cell populations grown in the other treatments display changes in proliferation and morphology as early as 6 hours (Figure 2a). For example, TGFB+EGF ligand treatment increased cell spacing and induced large lamellopodia, while OSM treatment induced tightly packed cell clusters; see Gross et al.^37^ for further phenotypic assessment. These changes motivate our interest in quantitative analysis of morphodynamical trajectories.

We characterized changes to cell morphodynamics under the different ligand treatments by quantifying the similarity between distributions of morphodynamical trajectories between ligand conditions (Figure 2b-d). We found similar ligand-specific distributions in the embedding space of morphological snapshots, with increased ligand-specific uniqueness observed in the embedding space of morphological trajectories. At the snapshot level, which excludes trajectory information, Figure 2b shows that over the course of the experiment cells occupy broad distributions in the embedding space. For example, we observed distinct shifts in occupancy that separate OSM and TGFB+EGF from other conditions, which is consistent with the distinct morphologies associated with these two treatments (Figure 2a). At a trajectory length of 8 steps (3.5 hours), these broader relationships are preserved but the cell state distributions in the embedded space become more condensed and display distinct peaks (Figure 2c). The uniqueness of cell state distributions between ligand treatments is reflected in a monotonic reduction in the shared area, or overlap, between cell state probability distributions with increasing trajectory length (Figure 2d). The pairwise overlap decreased more rapidly than in a null model in which cell features were randomly scrambled within treatment (Figure 2d). Thus, the trajectory embedding that leveraged information across timepoints yielded improved description of the ligand-specific morphodynamical responses as compared to snapshot analysis.

### Improved cell state description from morphodynamical trajectories

Cell states can be defined by identifying metastable regions of the morphodynamical trajectory embedding space where trajectories remain localized for extended time periods. To compare cell states and their relationships, we extracted the single-cell dynamics in the embedding space, i.e., the space of trajectory snippets defined in a sliding window. If there are *T* timepoints in a full single-cell trajectory, then there are *T-n_τ_* + 1 snippets of length *n_τ_*, each of which is a point in the embedding space. Together, these points trace out a trajectory in the embedding space with *T-n_τ_* transitions between snippets (e.g., from the snippet consisting of frames 1-8 to the snippet consisting of frames 2-9). We used the dynamical information about snippet-to-snippet transitions to calculate dynamics in the morphodynamical cell state space. The average of all cell state trajectories passing through a local region in the landscape yields the cell state “flow”, which in a Markovian picture of a continuous stochastic process^41, 42^, is proportional to the effective force “pushing” a cell from one morphodynamical state to another. These cell state force-fields are visualized in Figure 3. In the snapshot landscape, cell state trajectories appear highly random with little systematic variation between treatments (Figure 3, left column). In the trajectory embedding landscape, however, the cell state force-field displays treatment-specific convective flows (Figure 3, center column). These flows indicate stabilization in the unique regions of density peaks between treatments, providing direct evidence for the paradigm of metastable attractors in a landscape of cell state. Individual single-cell trajectories in the embedded space can stay partly localized to these metastable cell states for extended time periods. Over the timescale of 10+ hours, cell state changes reveal the transition pathways between metastable cell states (Figure 3, single-cell trajectories shown as blue to green lines, image sequences in the right column). Trajectories that appear random when observed via single-timepoint snapshot features unfold^43, 44^ and become systematic in the trajectory embedding space.

**Figure 3:**
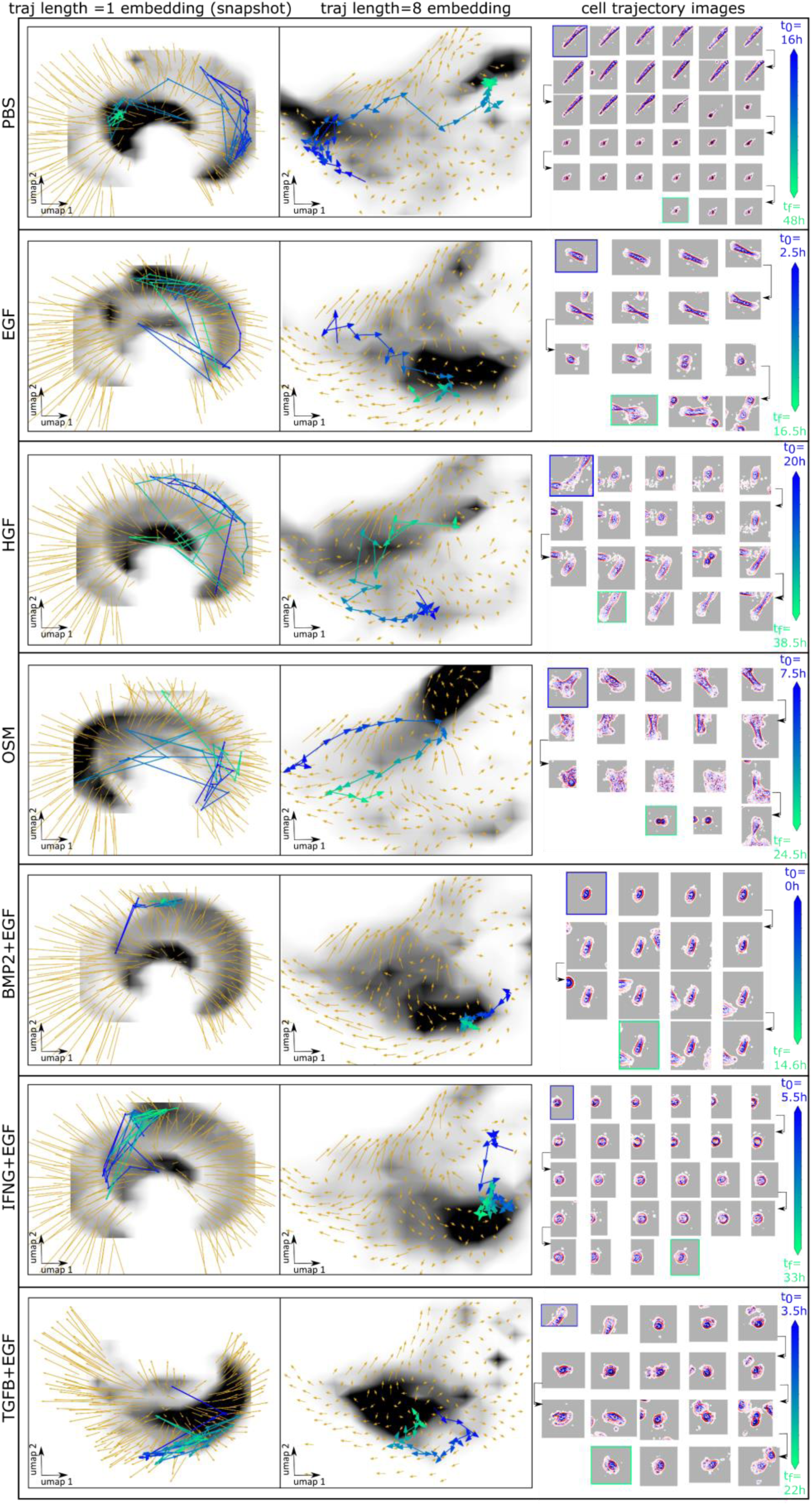
Trajectory embedding enables the determination of metastable cell states and pathways across ligand treatments. Left and middle column: average displacement proportional to the effective force (orange arrows), and cell density (grayscale) for snapshot embedding (left: snapshot, trajectory snippet length = 1, right: trajectory snippet length = 8). Representative cell morphodynamical trajectories capturing cell state transitions (blue to green line with arrows showing the direction of motion in the embedding space) from t_0_ to t_f_ determined by the available cell tracks. Right column: Cell images every hour along the extracted trajectory, with the tracked cell centered in the image frame and neighboring cells moving in and out of view, except in the IFNG+EGF images where the tracked cell is temporarily clipped at the edge of the microscope field of view.

The utility of a single-cell trajectory depends upon how well it characterizes cell state transitions and transition dynamics. We measured how systematic and predictable the trajectories are by quantifying the randomness of the trajectories in the embedding space. We defined the predictability of a trajectory as a locality ratio 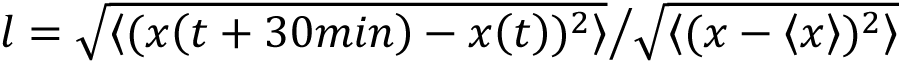 between the root-mean-square (RMS) displacement in the embedding space after one timestep (30 min) and the standard deviation in the displacement over the full population. Angled brackets 〈⋯ 〉 indicate averages over all trajectories and timepoints *t*. In a completely random trajectory, this ratio is 1 because the variance in a single timestep and the full population is identical. In a deterministic trajectory, all trajectories emanating from the same point are identical and have no variance after a single timestep--the only contribution to the ratio is the relative average displacement in the time interval. In a continuous, stochastic description of the trajectories in the embedded space^41, 42^, this locality ratio is related to the effective diffusion rate. Figure 4a shows that this locality ratio systematically decreases with trajectory embedding length in contrast with the null model, indicating that trajectories are increasingly less random and more predictable with increasing trajectory embedding length.

**Figure 4:**
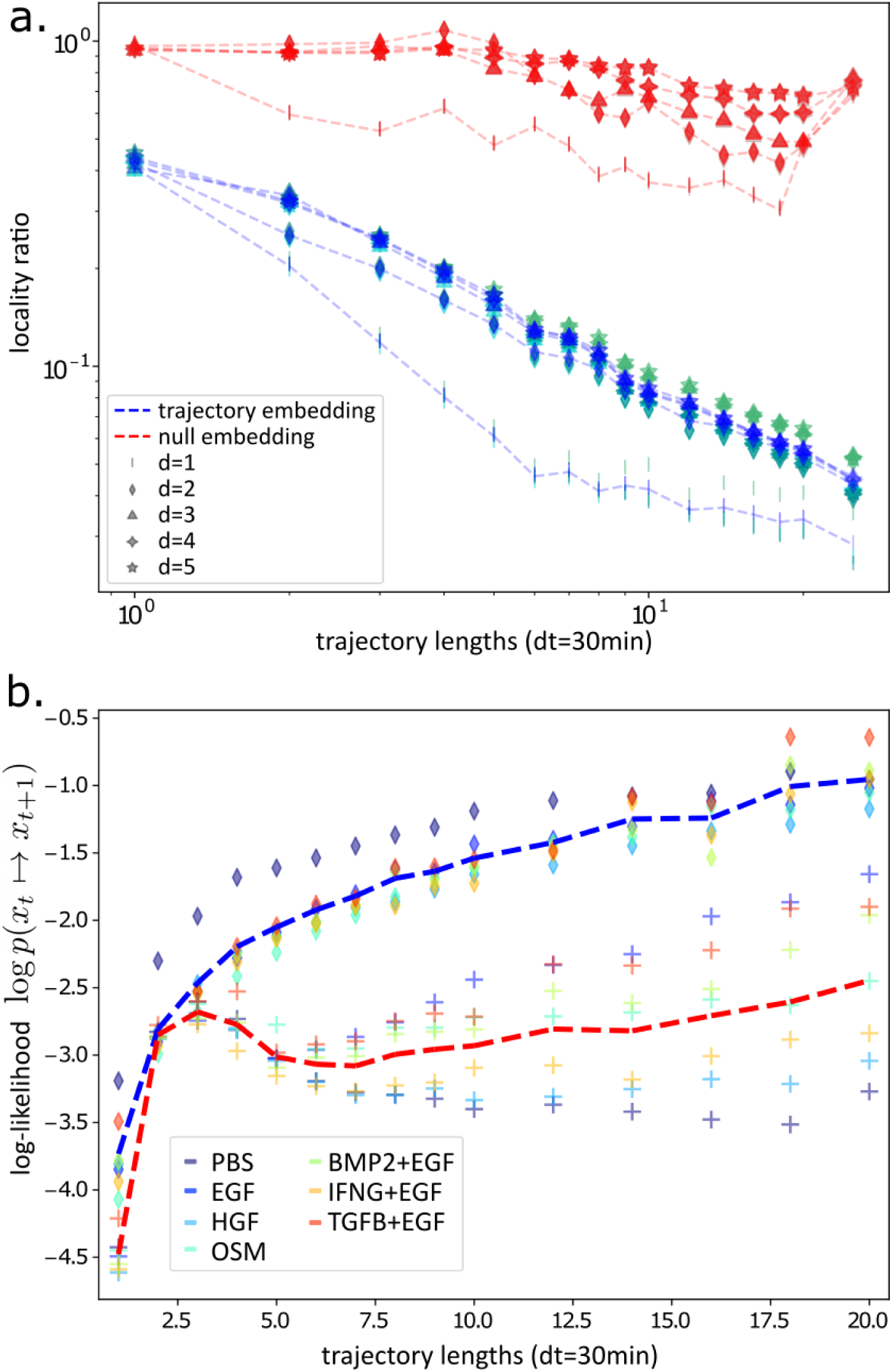
Trajectory embedding increases the predictability of cell trajectories. a) Ratio between the single-step (dt=30min) and full RMS displacement in the trajectory embedding space as a function of the trajectory length, null model with randomly scrambled features within treatments (reds) and trajectory embedding (blues), and UMAP embeddings with d=1 (lines), d=2 (diamonds), d=3 (triangles), d=4 (squares), and d=5 (stars), with d the number of UMAP components. Three replicates are shown per embedding (sea green, turquoise, teal). b) Average log-likelihood per trajectory step from the validation set cell trajectories, as a function of the trajectory length, averages for the trajectory embedding (blue dashed line) and for the null model (red dashed line), from UMAP d=2 embeddings. Individual treatments (colors) for the null model (crosses) and for the trajectory embedding (diamonds).

To determine the capability of the morphodynamical embeddings to characterize single-cell morphodynamical trajectories, we first used a subset of the data to train a model of trajectory likelihood, then calculated the average log-likelihood of a held-out test set of trajectories. The average log-likelihood is a direct measure of the predictability of the cell trajectories^45^. Here we used a Markovian transition matrix likelihood model, trained by counting transitions between Voronoi states defined by k-means cluster centers on the landscape^46, 47^. We utilized 100 k-means centers to discretize the morphodynamical embedding space, which provided sufficient Voronoi centers to capture the observed patterns of cell state flow between metastable cell states while still retaining adequate sampling of state-state transitions. The average log-likelihood increases as a function of morphodynamical trajectory embedding length and is higher than in a null model where cell features were randomly scrambled between treatments, shown in Figure 4b.

We expect in general that greater trajectory embedding lengths will increase the descriptive capability of trajectory models, but only up to the point where the increase in information in the longer time-ordered trajectories is greater than the loss of information due to incomplete tracking of cells between frames. For snippet lengths longer than 10 frames, we did not observe an increase in ligand-specific trajectories relative to the null model (Figure 2d), which likely is related to the decrease in the number of extracted trajectories longer than 8 frames (supplementary data table 2). Thus we chose a trajectory embedding length of 8 (3.5 hrs) for further analysis. The decreased overlap between ligand-specific distributions (Figure 2d), decreased locality ratio (Figure 4a), and increased trajectory likelihood (Figure 4b) indicate that even partial trajectory information over a timescale of a few hours substantially improved the representation of cell state.

### Morphodynamical transitions precede cell cluster formation

Ligand treatments displayed characteristic transitions between cell states (Figure 5a), indicating ligand-specific regulation of cell morphodynamics. In general, cell state distributions were more similar across ligand treatments at early times and became more condensed and distinct at later times, which is reflected in the time-dependent morphodynamical cell state distribution (supplementary figure 3).

**Figure 5:**
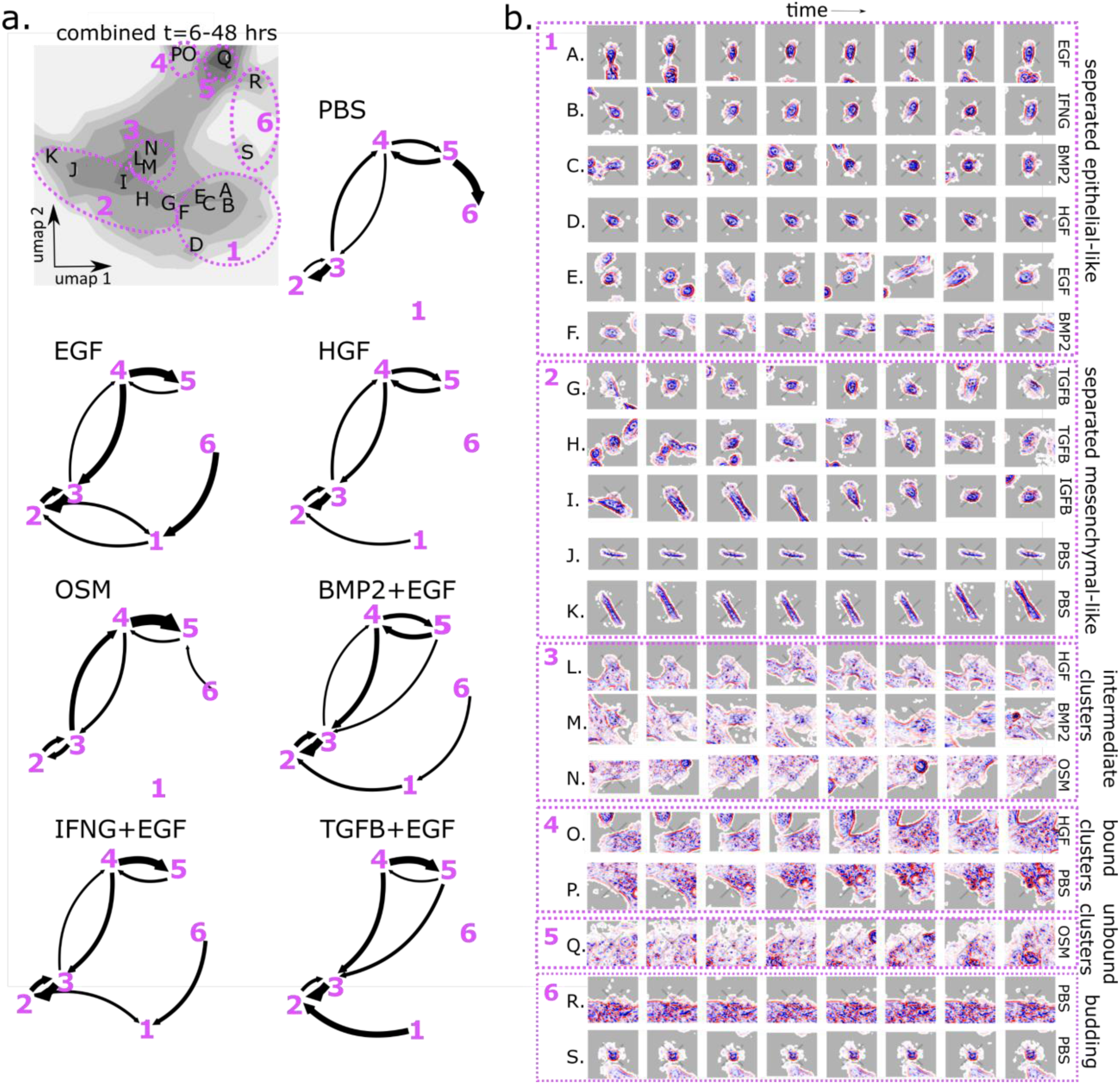
Trajectory embedding resolves pathway of cell cluster formation via mesenchymal-like intermediate. a) Cumulative distribution of all cells under all treatments (grayscale) with labeled fine-grained density peaks (A-S), and qualitative cell states (mauve dashed circles) with numeric labels (1-6), and cell state transition networks with arrow weight proportional to conditional transition probability, not overall transition flows, with transition probabilities <3% not drawn. b) Representative trajectory snippets (embedding length = 8 = 3.5 hrs) extracted at density peak locations (A.-S.) with the treatment condition of the representative trajectory snippet, grouped by macrostate with morphologically descriptive macrostate labels (right of images).

Cell states in the embedding space were identified using quantitative methods and refined by visual inspection. We identified fine-grained metastable states spanning the cell state landscape from density peaks arising in individual treatments (Figure 5a, 5b A.-S.), which we divided into 6 states, or groupings, and assigned qualitatively descriptive labels based upon observed cell morphology. These labels are intended to be descriptive for ease of interpretation. “Separated epithelial-like” cell states are rounder cells enriched in EGF condition, while “separated mesenchymal-like” cell states display more extended cytoskeletal features such as lamellopodia and are enriched in the TGFB condition. The “intermediate clusters” cell state represents cells that are mostly separated from tightly clustered cells. We further divided multicellular clustered cell states into “bound clusters” which display a thick border around attached cells that span multiple cells, “unbound clusters” which lack this border and whose outer cells have extended cytoskeletal features, and “budding” cells which consist of semi-round cells attached to the outer border of a cell cluster.

The direction of the cell state flow in the trajectory embedding space indicates that a group of cells coming together and forming an attached multicellular cluster proceeds via state 2, “separated mesenchymal-like” cells with extended cytoskeletal features. This flow was consistently observed in the cell state “force-fields” (Figure 4), cell state transition networks (Figure 5a), and the time-dependent cell state distributions (Supplementary Figure 3), and in all of the ligand conditions.

Cell cluster formation was most strongly associated with OSM treatment,^37^ consistent with figure 5a showing that in the OSM treatment the transition probability from the “intermediate cluster” state 3 into the multicellular clustered state was higher than the transition probability out of it. Separated cells (states 1 and 2) are separated from cell clusters (states 4 and 5) and cell cluster formation proceeds through states 2→3→4 via cells displaying extended cytoskeletal features and increased cell border contrast (Figure 5b).

## Discussion

In vivo, cells continually modulate their phenotypic state in response to changes in local microenvironmental cues^38^. During development, cells must precisely control cell states^48^, responses to extracellular signals, and cell motility, and the loss of cell state control is associated with various diseases, including cancer^49^. Dynamic cell behaviors can be observed via live-cell imaging but quantifying the dynamic relationships between diverse cellular phenotypes has been challenging. Our morphodynamical “trajectory embedding” provides a method to quantitate dynamic morphologic behaviors. Trajectory embedding leverages the unique capability of live-cell imaging to follow single cells in time and constructs a coordinate space based on time-aggregated “hyper features” better suited to study cell states and their dynamical relationships as compared to standard featurization based on single timepoints.

We observed that cells dynamically transition between morphodynamical cell states and that the transition frequency is strongly modulated by ligand treatment. These dynamic cell state relationships can provide a framework for understanding cell-cell heterogeneity and heterogenous cell responses to perturbation. Live-cell trajectory embedding brings the cell state landscape paradigm from theoretical biology to direct application, where cell states and the transitions between them can be resolved, validated, and potentially leveraged for actionable control strategies^14, 50–55^.

Our morphodynamical trajectory embedding procedure quantifies the space of morphological trajectories directly, leading to an improved description of dynamical cell state changes compared to using only morphological snapshots. One limitation of our analysis is that we did not explicitly consider cell cycle stage, instead, these processes remain implicitly described via morphological features. We envision that future studies could extend our framework by directly identifying cell cycle stage and mitotis. Such an extension would provide insight into the coupling of cell cycling to changes in motility and morphology and the heritability of morphodynamical state from parent to daughter cell. We chose a broad cell feature set but many other approaches to define cell features have been developed, including other novel cell shape descriptors^56, 57^ and machine learning-based approaches^58^. Trajectory embedding, in principle, has the capability to map dynamical information from any reasonable featurization towards a more complete description of cell state.

The morphodynamical trajectory embedding method can be applied to any live-cell imaging modality where cells can be characterized and tracked through time, and in particular, we expect that this method will be especially powerful for analysis of live-cell approaches that incorporate genetically-encoded, fluorescently-labeled reporters, both *in vitro*^59–61^ and *in vivo*^62–64^. Practical limitations will always come into play, nevertheless. For example, we applied our method to unlabeled phase-contrast imaging of cell cultures that approach confluence^65, 66^, without paired ground truth labeled images^67^, which necessitates an analysis approach with some robustness to cell segmentation and tracking errors. Trajectory data quantity and quality will generally pose a constraint on trajectory length used in the morphodynamical trajectory embedding analysis. Even when many very long trajectories are available, analysis based upon shorter trajectory lengths might more directly capture processes and relationships of specific interest, such as the connection between specific morphological features and cell motility. General procedures to identify the optimal trajectory length for analysis need to be developed, building on the preliminary approaches employed here.

Biological interpretation and validation of the morphodynamical cell states extracted here will be important, and our findings help to motivate specific hypotheses that could be explored in future studies. Identification of the molecular programs associated with particular cell states and cell state transitions would provide insight into how these processes are mediated in normal tissues and how they may go awry in diseases. Importantly, trajectory embedding analysis enables quantification of cell state transitions, and may therefore be useful for gaining insights into disease progression and therapeutic resistance^68–74^. Our observation that cell cluster formation is preceded by a mesenchymal-like shift in cell state aligns with the maturation of transverse arc stress fibers as a precursor to stable cell-cell junctions observed by Rajakylä et al,^75^ but live-cell imaging coupled with deeper molecular profiling data—such as multiplexed imaging^76–79^ and single-cell transcriptomics^80, 81^—are needed in order to develop the applicability and utility of the information obtained from live-cell imaging alone. Manifold-based or mutual-information approaches have had some success with single-cell data integration^82–84^, and may enable integration of live-cell imaging trajectory embeddings with molecularly resolved data, a critical data analysis goal needed to provide insight into the biological relevance of morphodynamical cell states. With the trajectory embedding method we present here, we can now study the emergence of metastable attractors and the regulation of dynamic cell state changes, directly confirmed via live-cell trajectories.

## Methods

### Live-cell imaging of MCF10A cells

Data used in this study were recently described by Gross, et al.^37^ In brief, MCF10A cells were plated at 75,000 cells/well on collagen-coated 8-well plates. After an 8 hour attachment period in growth media containing EGF and insulin and a 12 hour period in media lacking EGF and insulin, cells were treated with 7 different ligand treatments (PBS, EGF 10 ng/ml, HGF 40 ng/ml, OSM 10 ng/ml, BMP2 20 ng/ml + EGF 10 ng/ml, IFNG 20 ng/ml + EGF 10 ng/ml, TGFB 10 ng/ml + EGF 10 ng/ml). Wells were imaged every 30 minutes for 48 hours via bright-field phase contrast with an Incucyte microscope (1020×1280, 1.49 *μm*/pixel), with the initial frame coinciding with the addition of the ligands and fresh imaging media. 6 image stacks were collected for each ligand treatment and imaged simultaneously, see Gross et al. ^37^ for a discussion of batch effects and experimental replicates. Experimental protocols can be found in detail at the publicly available Synapse database^85^.

### Image preprocessing

Foreground (cells) and background pixel classification was performed using manually trained random forest classifiers using the ilastik v1.3.3 software^86^. Images were z-normalized (mean subtracted and normalized by standard deviation) and background pixel values were set to a value of 0. In cell images, these z-normalized pixel values are shown from red to blue (positive to negative).

### Cell segmentation

Single-cells were segmented from the preprocessed images using the cytoplasm model of the Cellpose^87^ v0.6.5 software, a deep learning approach trained broadly across cell types and imaging modalities. The Cellpose algorithm requires estimating the size of the cells before segmentation; due to the variability in sizes and shapes of the MCF10A cells, segmentation was performed iteratively over multiple rounds allowing the Cellpose algorithm to determine a new cell size at each round until no more new cells were found (pixels of previously segmented cells set at each round to the background value of 0). Image preprocessing and segmentation scripts can be found on the github repository, see data and code availability. The unlabeled, bright-field phase-contrast imaging used here leads to image analysis challenges, particularly for cell segmentation. It is difficult to judge the quality of many extracted cell segmentations when local cell density is high, see Supplementary Table 1 for manual validation.

### Cell featurization

Three classes of features were used to characterize individual cells: (i) texture features, (ii) shape features, and (iii) features characterizing adjacent cells. As preliminary steps, segmented cells were extracted, and mask-centered into zero-padded equal sized arrays larger than the linear dimension of the biggest cell (in each treatment); then the long axis was defined by the non-mass-weighted moment of inertia of the cell mask and aligned along a reference axis. (i) Two types of internal cell features were calculated. Zernike moments (49 features) were used to characterize the overall spatial phase contrast signal and Haralick texture features (13 features) were used to characterize the phase contrast texture; these were calculated in the Mahotas^88^ image analysis package. The sum average Haralick texture feature was discarded due to normalization concerns. (ii) Shape features (15 features) were calculated as the absolute value of the frequency coefficients of the Fourier transform of the distance to the boundary as a function of the radial angle around cell center^89^, with the sum of shape features normalized to 1. (iii) The cell environment was featurized in a similar fashion, where an indicator function with value 0 if the cell boundary was in contact with the background mask (no neighboring cell), and value 1 if in contact with the cell foreground mask. The absolute values of the Fourier transform coefficient of this indicator as a function of radial angle around the cell-center then featurized the local cell environment (15 features), with the sum of cell environment features normalized to 1. Note the first component of the cell environment features is practically the fraction of the cell boundary in cell-cell contact. Additional information regarding the cell featurization can be found in Supplementary Figure 4A.

After computing the raw features as described, the high-dimensional cell feature space was dimensionally reduced using principal component analysis (PCA), retaining the largest 3 eigen-components of the feature covariance matrix (spanning all treatments and image stacks) which captured >90% of the variability. More PCA components can be retained with only a small computational cost in the dimensionality reduction step (here UMAP) which typically scales linearly with the data vector length. Additional information regarding the cell feature PCA reduction can be found in Supplementary Figure 4B.

### Cell tracking

Segmented cells were tracked between live-cell imaging frames to extract the set of single-cell trajectories. Image stacks were first registered translationally without allowing rotational or general affine transformation using the pystackreg implementation of subpixel registration^90^. Cell centers were recorded as the equally weighted center of mass of the single-cell masks. Cells were tracked between frames by first separating each contiguous cluster of cells as defined by connected sets of the foreground/background cell mask. If cell clusters occupied less than 10,000 pixels (typical cell very roughly 30×30 pixels), cells were simply tracked by minimum distance with a cutoff of 45 pixels. Tracking cells by minimum distance refers to linking a cell at frame t to the cell at frame t+1 which has the minimum distance between cell centers. For larger cell clusters, clusters were first tracked by minimum distance with a cutoff of 300 pixels. Tracked cell clusters were each individually registered rotationally and translationally (again using pystackreg), and individual cells in the clusters were tracked between frames by maximum overlap with a cutoff of 10 pixels. The ligand panel yields differential impacts on cell proliferation, with a typical cell division time of ∼12 - 20 hours. In a correctly tracked cell division event, a parent cell will be linked to two daughter cells, which in turn will lead to two separate trajectories sharing a common initial history prior to mitosis. In our analysis, two or more cells may have the same parent cell as the closest cell in the previous frame. This can be the result of a cell division event, or can be the result of a missed track or missed cell segmentation. We do not separate cell division events into their own category, but simply use the cell tracks, or linkages between frames, to identify the unique cell trajectory history. Where trajectories are split as in a cell division, trajectories are not truncated and begun anew, but rather each daughter is tracked backward in time leading to two trajectories which are treated separately, see cell image sequences from HGF and TGFB+EGF conditions in Figure 3 for examples. That is, daughter cells will have overlapping or shared history from the parent cell.

### Morphodynamical trajectory embedding

High-dimensional time-sequences of features are used as the input to dimensionality reduction algorithms as the basis for morphodynamical cell state analysis. Extracted cell linkages were constructed from available cell tracks. These cell linkages are not complete, that is incomplete segmentation/tracking means that some cells cannot be traced all the way back to an initially plated progenitor cell. The available cell history is the unique backwards trace of any extracted single-cell through time. Of the total 476,855 cells extracted here; 137,845 could be traced back only one step, while 36,919 could be traced back 10 steps, see Supplementary Table 2. We consider the set of all cells (from possibly different experimental time points) with cell history equal to or longer than a given length, the trajectory snippet set, see Supplementary Figure 5 for a graphical description. Note there is a large amount of duplication in this sliding window division of available cell trajectories. Cell trajectories that are longer than the trajectory snippet length used in the trajectory embedding allow for the determination of cell states and pathways. From the linkages, all cell trajectory snippets of length *n_τ_* (all possible cell histories with length *n_τ_*) were extracted in a sliding window manner. The number of available trajectory snippets for each treatment is shown in Supplementary Table 2. The PCs of each cell in the trajectory snippet were then concatenated together (e.g. for 2 PCs 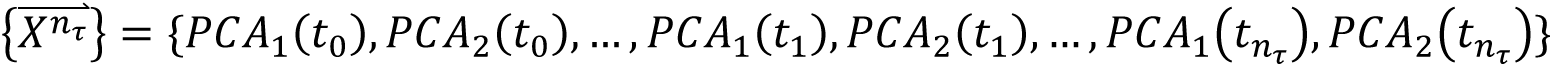 to form the trajectory snippet supervector, which we define as the morphodynamical feature trajectory of length *n_τ_* (with N features, the feature trajectory for each cell is N x *n_τ_*). These morphodynamical feature trajectories (spanning all image stacks and all treatments) are flattened into vectors and then embedded using UMAP^91^ into a space of dimension d. Changing UMAP embedding hyperparameters will alter fine details regarding the trajectory embedding landscape, but we find our overall results to be robust with very little change in the measured overlap between ligand populations or trajectory likelihood as shown in supplementary figure 6. The trajectory embedding analysis allows for the robust and systematic characterization of cell state trajectories even in this challenging data analysis regime with many missing and partially segmented cells.

### Overlap coefficient

To compare the similarity of two probability distributions over a shared space, we use the overlap coefficient^92^ defined by the sum of the minimum value of two probability distributions. The overlap is 0 for completely distinct non-overlapping distributions, and 1 for identical distributions.

### Stochastic dynamics: locality ratio, cell state force-fields

We employ several measures motivated by standard concepts of stochastic physics. A measure of the randomness of motion in stochastic dynamics is the effective diffusion rate defined at a timescale *τ* by *D* ≡< Δ*x*^2^ >_*τ*_/*τ* with here *τ* the time between frames of 30 minutes. We characterize how random trajectories are by the ratio of the single-step RMS (root-mean-square) displacements to the total RMS displacement, a locality ratio 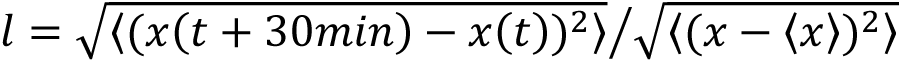 where the mean-squared displacements are summed over dimension and the angle brackets 〈… 〉 indicate averages over all trajectories and time points. For completely random trajectories, this ratio is 1, and tends to the relative average displacement for a continuous, deterministic dynamic. If cells are obeying stochastic Markovian dynamics characterized by a diffusion equation, then the average displacement 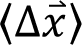 is proportional to the effective force 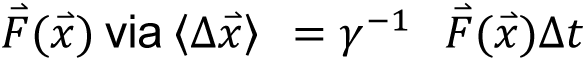 with *γ* the friction.

### Cell state clustering and prediction

The cell state dynamics were characterized by building a fine-grained discretized transition matrix model in the continuous morphodynamical trajectory embedding space. The embedded space was binned using k-means clustering with *k* = 100 clusters. In the discrete space, a transition matrix between bins was estimated from the observed transition counts *C_ij_* from a microbin *i* to microbin *j* as *T_ij_* = *C_ij_*/*C_i_* and *C_i_* = ∑_*j*_ *C_ij_*. This transition matrix is commonly referred to as a “Markov Model” broadly used in the analysis of molecular systems.^93^ Note that this transition matrix does not share a steady-state distribution with the cell populations, as cell birth and death states are not included. This transition matrix was used as a (highly simplified) model of the single-step trajectory likelihood.

### Trajectory likelihood

To assess the quality of the description of the cell state dynamics, we adopted a self-consistent measure of how likely a test set of single-cell trajectories were within a transition matrix model trained from a separate training set of trajectories. Data were split into a training set (5/6 images stacks per treatment) and a validation set (1/6 image stacks per treatment). The training set was used to train a transition matrix likelihood model, and the average log-likelihood per trajectory step was calculated from the test set trajectories using the transition matrix as 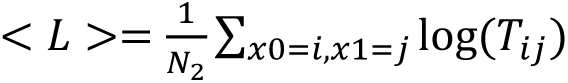 with *N*_2_ the set of all 2-step trajectories in the validation set (initial point *x*0 mapped to bin *i* and next point *x*1 mapped to bin *j*.

### Cell metastable state extraction and grouping

To capture the broad relationships defined in the continuous trajectory embedding space we defined a small set of discrete cell “states”. We first picked out all metastable locations on the landscape. We identified these metastable locations by local maxima in the population density, see Figure 5 and supplementary figure 3. These fine-grained metastable cell states spanning the trajectory embedding landscape (19 total) were picked via density peaks in the individual treatments. To group these fine-grained cell states into broader state groupings, we first utilized an unsupervised kinetic clustering approach^94^ which separated three major metastable basins consistent with the regions of the landscape enriched in the EGF, TGFB+EGF, and OSM conditions (respectively lower right, lower left, and upper regions, Figures 2B and 5C). We then manually refined these regions to distinguish between clustered cell states (states 4, 5, and 6) differentially occupied between ligand treatment. These 6 qualitative cell states were used to define cell state transition networks (transition matrix, see **Cell state clustering and prediction** and Figure 5a). Cells were mapped into these states by first finding the closest fine-grained metastable state in the embedding space, and then assigning the state label accordingly. The morphodynamical trajectory embedding space indicates a continuum of cell states, thus intermediate metastable states as indicated by density peaks which are nearest-neighbors but assigned to different states, such as E and F assigned to the epithelial-like state and G and H assigned to the mesenchymal-like state, may be very similar and not strictly distinct. Cell state names are descriptive for ease of interpretation but not based upon validated biological interpretation.

### Statistics and Reproducibility

Overlap and locality ratio results were validated by calculating over 3 replicates of the data, split into 3 groups composed of 2 image stacks for each treatment. Means over the replicates, and the individual replicate data points are plotted to allow visual estimation of the robustness of the analysis. Error bars and 95% confidence intervals are estimated from the data splits by Bayesian bootstrapping^95^.

## Acknowledgments

We thank Ian McLean and Mark Dane for project guidance and assistance accessing LINCS data, David Aristoff and Gideon Simpson for mathematical insights, and John Russo and Luke Ternes for input regarding computational implementation. J.C. is supported by the Damon Runyon Cancer Research Foundation Quantitative Biology Fellowship DRQ-09-20. Y.H.C is supported in part by the National Cancer Institute (U54CA209988, U2CCA233280, U01 CA224012). D.M.Z. acknowledges support from the OHSU Center for Spatial Systems Biomedicine and NSF grant MCB 2119837. These studies were supported in part by NIH research grants U54-CA209988 and U54-HG008100, and the Anna Fuller Foundation to L.M.H.

## Author Contributions

Conceptual development: JC, YHC, LMH, DMZ. Data collection: JC, SG. Data analysis and computational implementation: JC. Manuscript writing: JC, YHC, LMH, DMZ.

## Data and Code Availability

All codes and scripts to perform the analysis in this work can be found in the project github repository. The current version is linked below and an archived release at the time of publication can be found at (DOI: 10.5281/zenodo.7644765), including a tutorial with instructions to access a portion of the live-cell imaging data available for download. The full live-cell imaging dataset is available from the authors upon request.

https://github.com/jcopperm/celltraj

LINCS MCF10A Molecular Deep Dive data is available in some formats from the synapse database^85^ and additional data is available upon request.

## Supplementary Figures and Tables

**Supplementary Data Table 1.** Segmentation and tracking manual validation. 100 cells per treatment were randomly selected, and evaluated by eye to qualitatively assess segmentation and tracking accuracy. Fraction segmented was estimated by the image area covered by segmented masks divided by the area selected as being occupied by cells from the ilastik random forest pixel classifier. *(+EGF)

| ligand | total | EGF | HGF | OSM | IFNG* | BMP2* | TGFB* |
| --- | --- | --- | --- | --- | --- | --- | --- |
| % segmented | 52% | 53% | 51% | 46% | 54% | 52% | 42% |
| % good seg | 43% | 52% | 38% | 33% | 48% | 53% | 56% |
| % bad seg | 26% | 26% | 24% | 7% | 25% | 33% | 34% |
| %ambiguous seg | 31% | 22% | 38% | 60% | 27% | 14% | 10% |
| % tracked | 60% | 51% | 56% | 55% | 60% | 58% | 79% |
| % good tracks | 87% | 95% | 91% | 42% | 94% | 96% | 95% |
| % bad tracks | 2% | 5% | 0% | 3% | 6% | 0% | 0% |
| %ambiguous tracks | 11% | 0% | 9% | 55% | 0% | 4% | 5% |

**Supplementary Figure 1.**
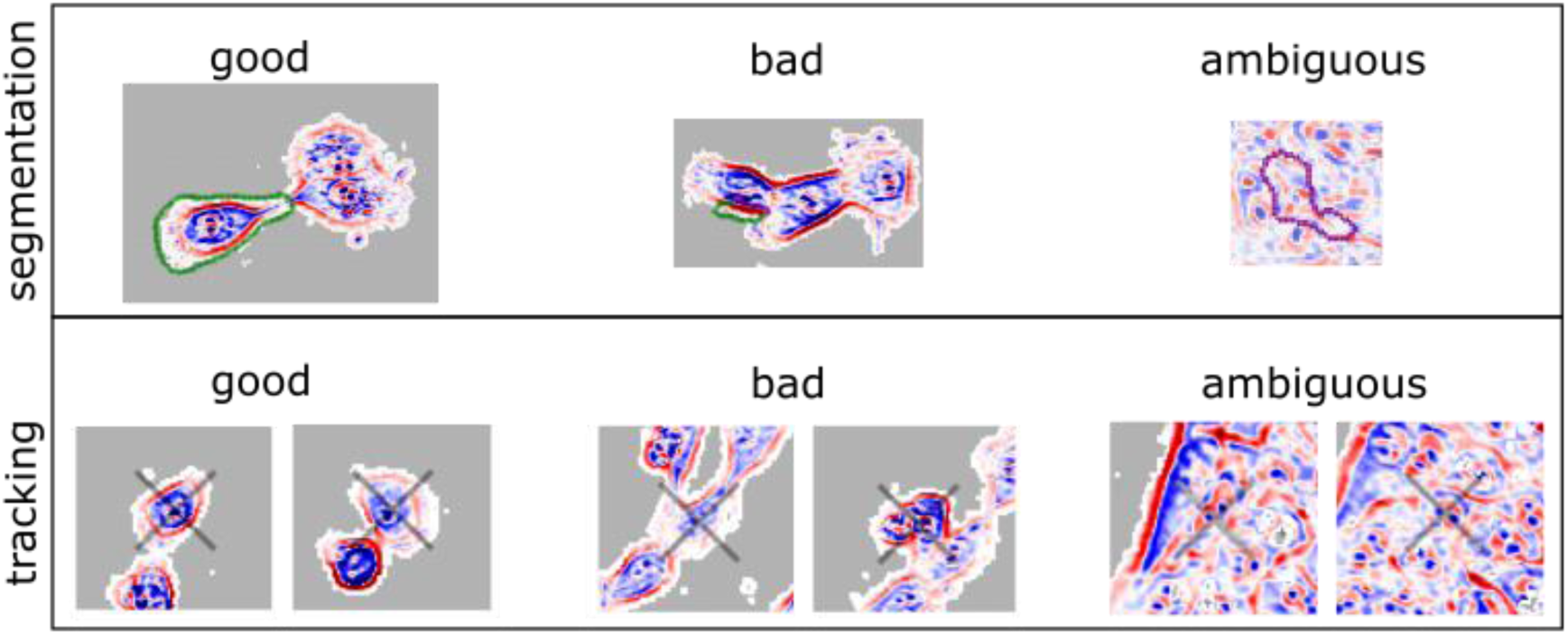
Segmentation and tracking manual validation. Examples of good, bad, and ambiguous qualitative validation categories for segmentation and tracking.

**Supplementary Data Table 2.** Number of extracted trajectory snippets with increasing snippet length. *(+EGF)

| Trajectory snippet length | PBS | EGF | HGF | OSM | BMP2* | IFNG* | TGFB* |
| --- | --- | --- | --- | --- | --- | --- | --- |
| 1 | 49016 | 73093 | 69747 | 99933 | 63669 | 72389 | 49008 |
| 2 | 32251 | 35337 | 40640 | 58119 | 31558 | 39545 | 36736 |
| 4 | 21894 | 14066 | 22837 | 29783 | 14135 | 19501 | 16206 |
| 8 | 15342 | 4892 | 12572 | 13208 | 5772 | 9033 | 2936 |
| 16 | 9256 | 1416 | 5857 | 3861 | 1892 | 3603 | 884 |
| 32 | 4243 | 226 | 2161 | 924 | 409 | 1055 | 142 |
| 64 | 1002 | 16 | 347 | 139 | 0 | 141 | 0 |

**Supplementary Figure 2:**
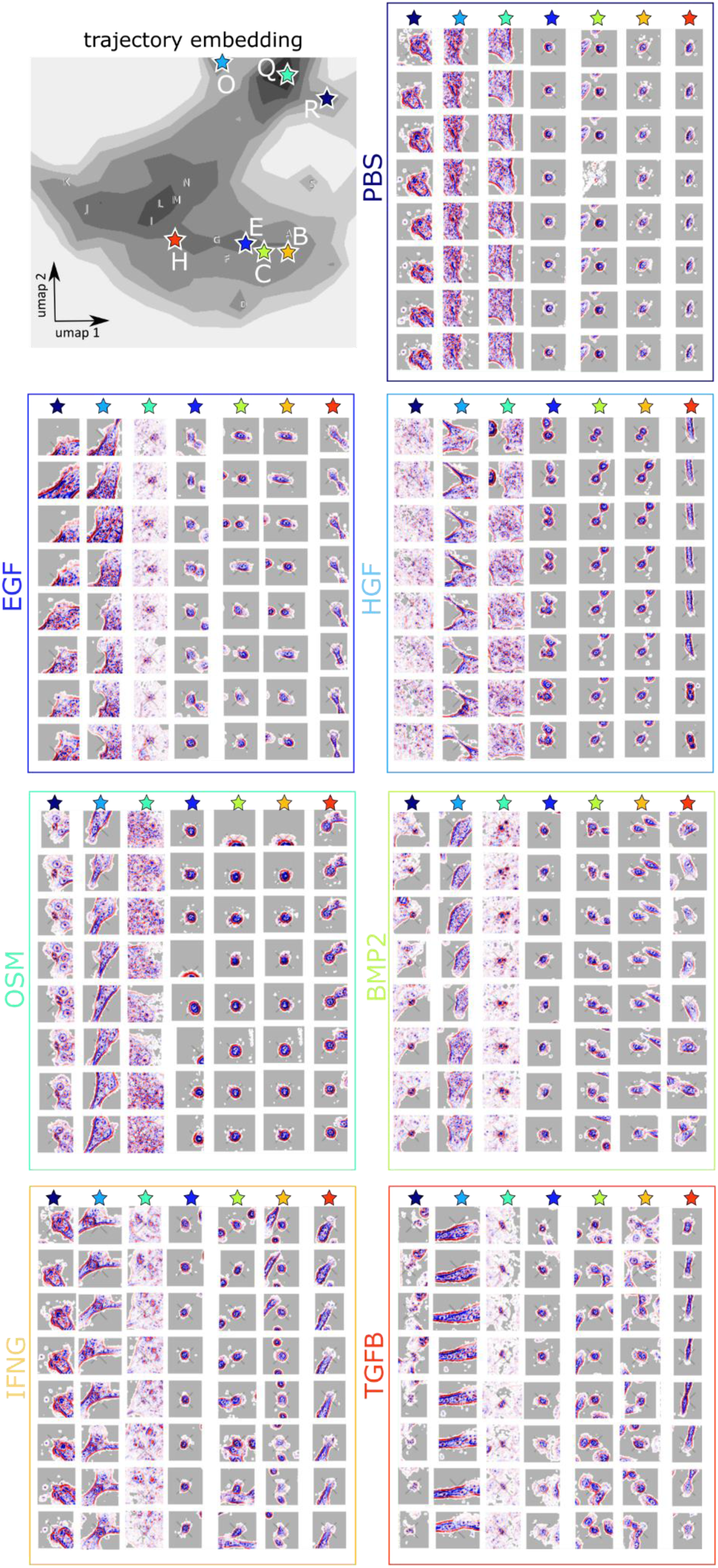
Trajectory embedding constructs a common space to evaluate unique and shared cell morphodynamics. Top left: outline of the combined density distribution in the trajectory embedding (snippet length = 8) space (gray), with locations of the density peaks in individual treatments marked with letters consistent with Figure 5 and exhibited cell trajectory snippets at locations marked with stars. Remaining boxes: cell trajectory snippet extracted at the marked location, but from the treatment labeled for each box. Time for each 8-step trajectory snippet shown runs from top to bottom.

**Supplementary Figure 3:**
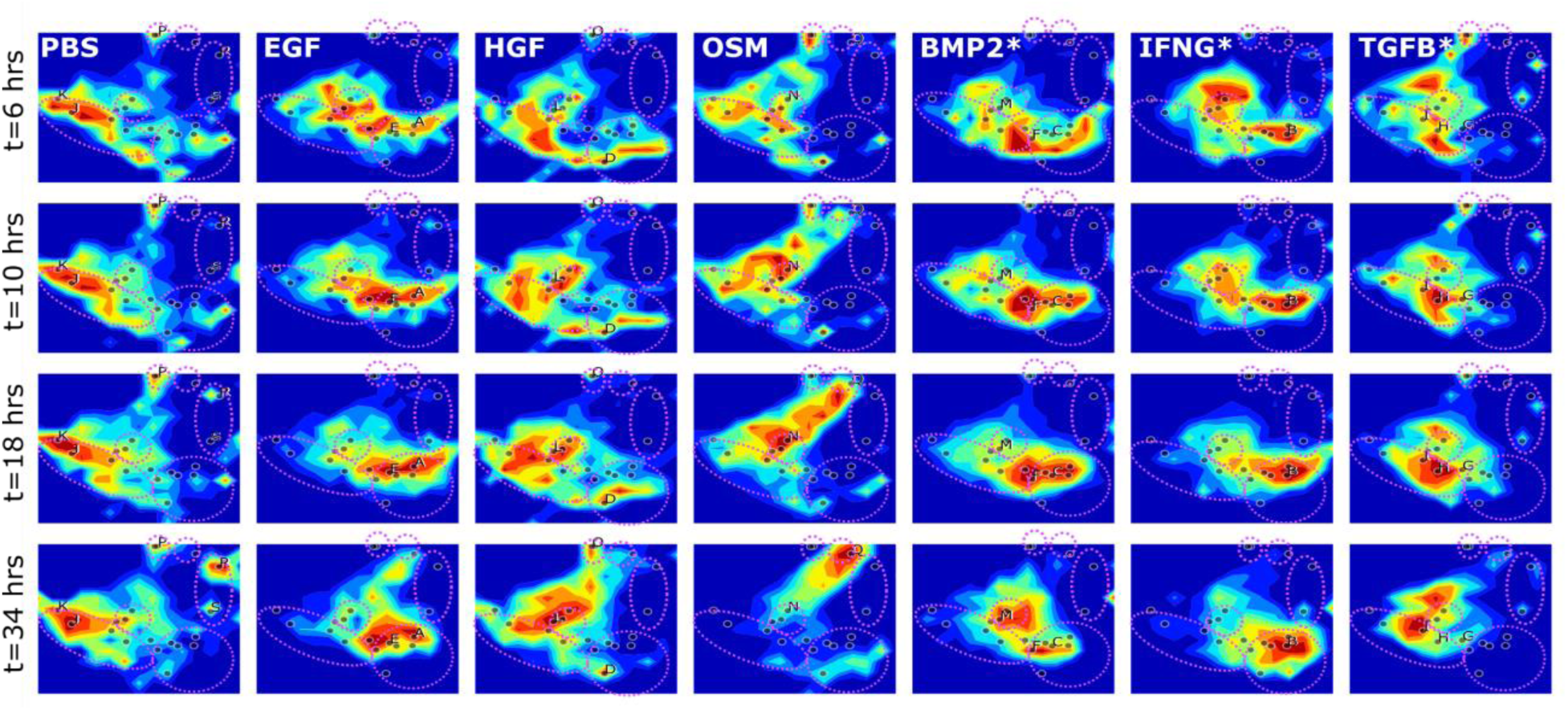
Time-dependent cumulative distributions (rainbow) from 12-hr windowed averages (or maximum allowed window average), and density peak locations selected as fine-grained metastable states (labels A-S). *(+EGF)

**Supplementary Figure 4:**
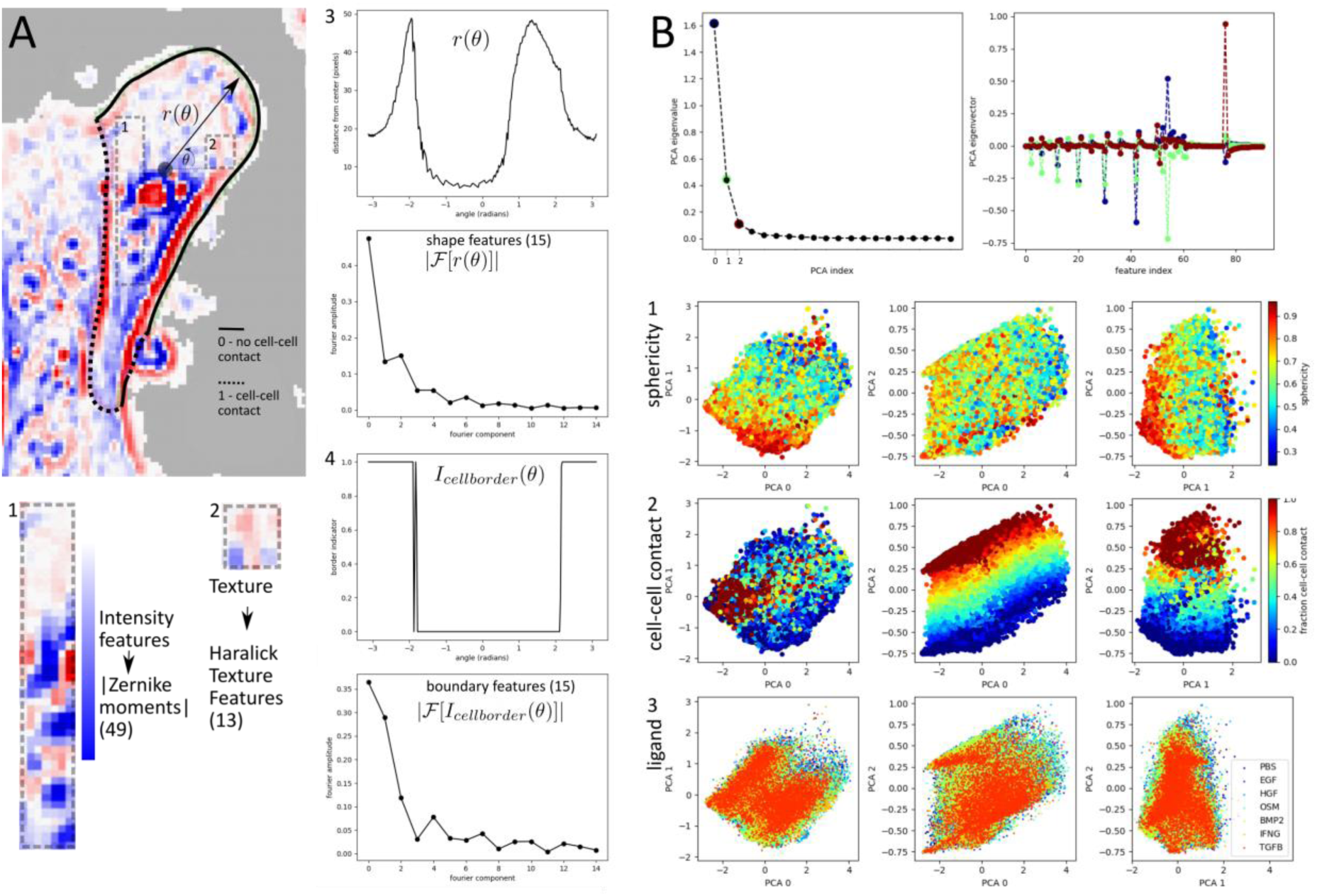
Cell features and PCA reduction. A) Segmented single-cell with cell border in contact with another cell (dashed line), and not in contact (solid line), and cell feature description 1. Phase contrast overall features described by Zernike moment absolute values. 2. Texture described by Haralick texture features. 3. Cell shape featurized by the absolute value of the Fourier coefficients of the distance to the cell center as a function of the angle *θ*. 4. Local cell environment described by the absolute value of the Fourier transform of *I*(*θ*) indicating cell-cell contact. B) PCA feature reduction. PCA eigenvalues (top left) and eigenvectors for the top 3 components explaining >90% of the feature variance (top right). PCA landscape colored by 1. Approximate sphericity given by the first Fourier component of *r*(*θ*), approximate fraction of the cell boundary in contact with another cell given by the first Fourier component of the cell-cell border indicator *I*(*θ*), and 3. by ligand condition.

**Supplementary Figure 5:**
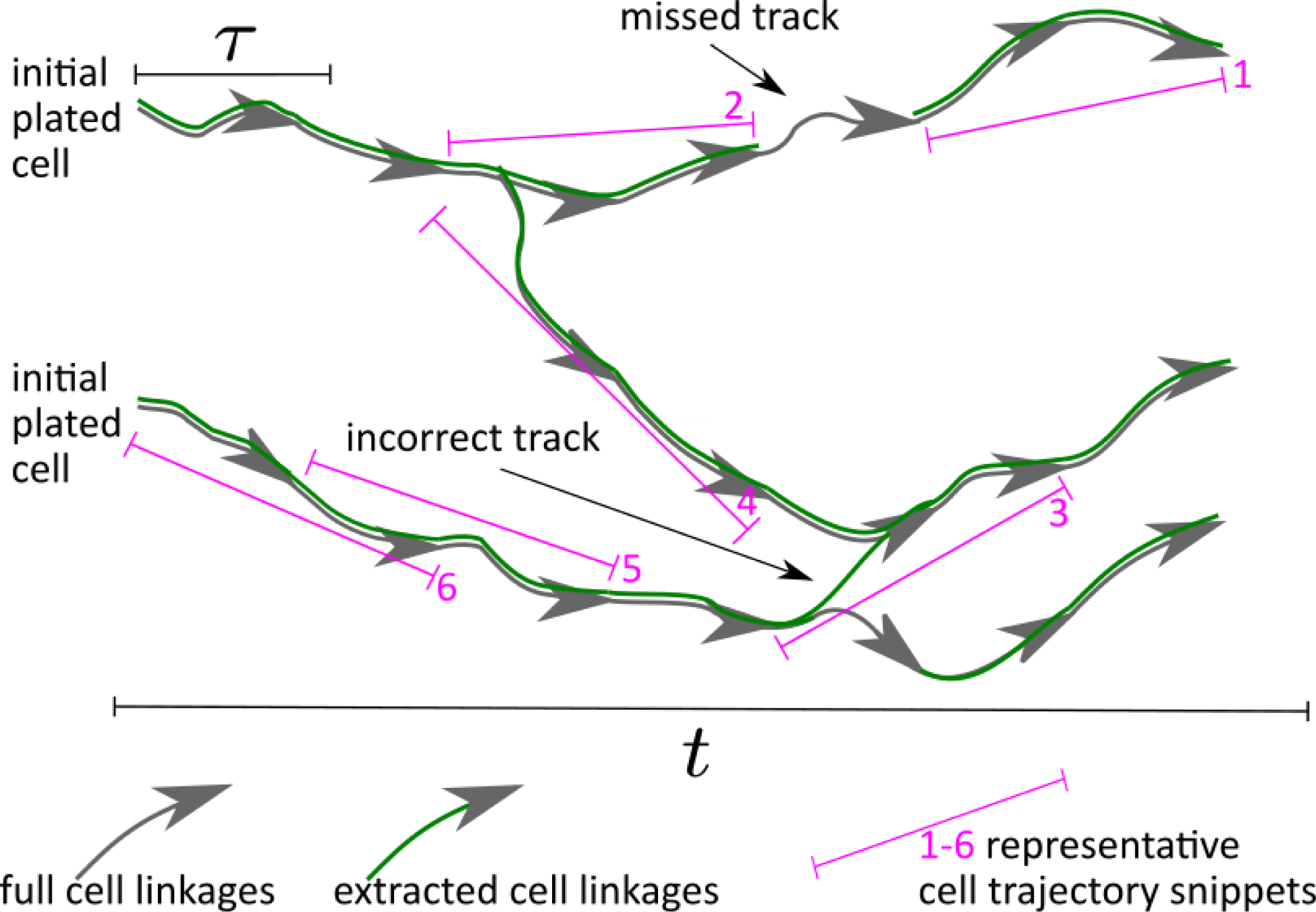
Graphical illustration of the full set of cell linkages connecting 2 initially plated cells to the cells at the final timepoint (gray arrows). The extracted cells and linkages from the cell segmentation and tracking steps as green arrows indicating the available partial set of cells and linkages used in the data analysis with errors and missing cells, and some possible trajectory snippets (yellow highlights) extracted in a sliding window manner along the extracted linkages.

**Supplementary Figure 6:**
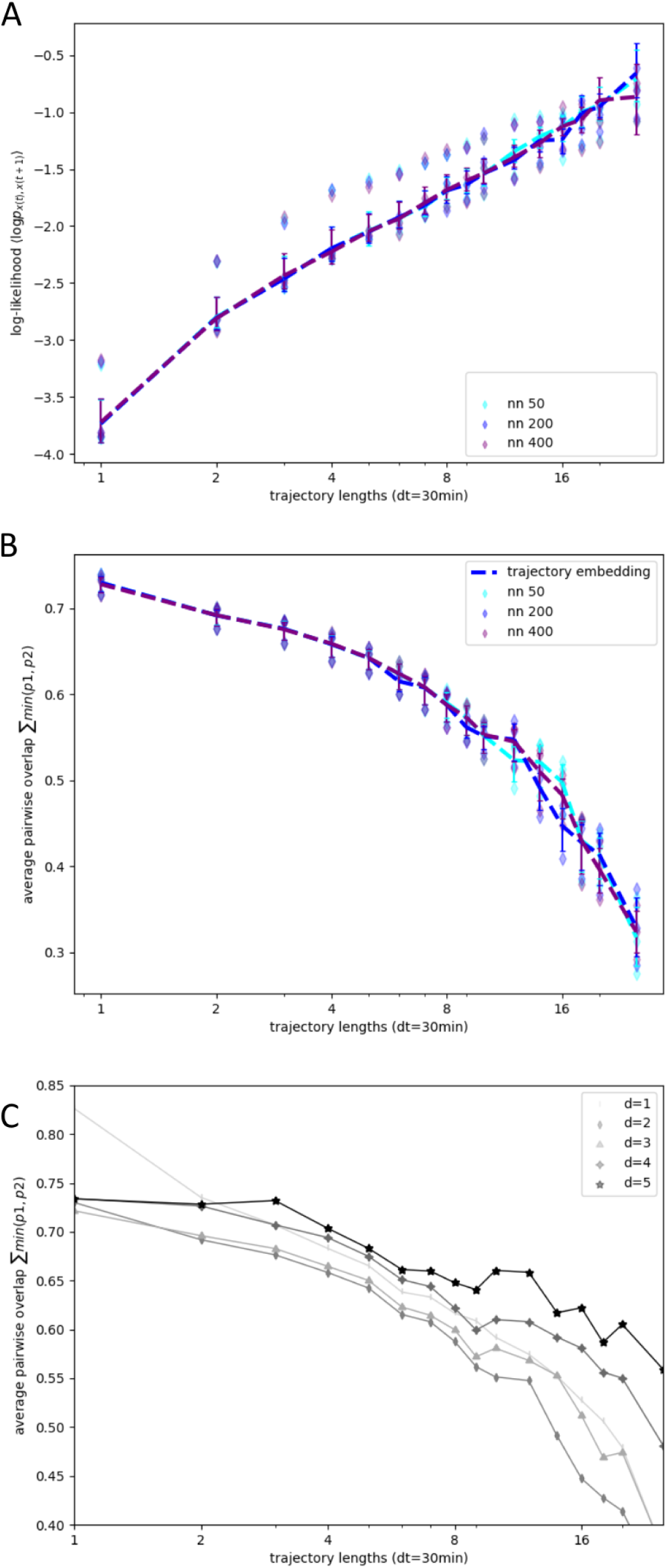
A) Trajectory log-likelihood with trajectory length at half the value of the UMAP n_neighbors parameter (nn=50, cyan) and twice the value of the n_neighbors parameter (nn=400, purple), and the value used in Figures 2-5 (nn=200). B) Average pairwise overlap between ligand population distributions at half the value of the UMAP n_neighbors parameter (nn=50, cyan) and twice the value of the n_neighbors parameter (nn=400, purple), and the value used in Figures 2-5 (nn=200). C) Average pairwise overlap between ligand population distributions with UMAP embedding dimension (d=1-5, light-gray to black).

